# Storage of prescription veterinary medicines on UK dairy farms: a cross-sectional study

**DOI:** 10.1101/342360

**Authors:** Gwen M. Rees, David C. Barrett, Henry J. Buller, Harriet L. Mills, Kristen K. Reyher

**Author notes:** Gwen Rees BVSc (Hons), MRCVS (Corresponding author,). Professor David Barrett BSc (Hons), BVSc (Hons), DBR, DCHP, Dip ECBHM, FHEA, FRCVS; Professor Henry Buller BA (Hons), PhD; Dr. Harriet Mills BSc, MRes, PhD; Dr. Kristen Reyher BSc, DVM, PhD, MRCVS.

## Abstract

Prescription veterinary medicine (PVM) use in the United Kingdom is an area of increasing focus for the veterinary profession. While many studies measure antimicrobial use on dairy farms, none report the quantity of antimicrobials stored on farms, nor the ways in which they are stored. The majority of PVM treatments occur in the absence of the prescribing veterinarian, yet there is an identifiable knowledge gap surrounding PVM use and farmer decision making. To provide an evidence base for future work on PVM use, data were collected from 27 dairy farms in England and Wales in Autumn 2016. The number of different PVM stored on farms ranged from 9-35, with antimicrobials being the most common therapeutic group stored. Injectable antimicrobials comprised the greatest weight of active ingredient found while intramammary antimicrobials were the most frequent unit of medicine stored. Antimicrobials classed by the European Medicines Agency as critically-important to human health were present on most farms, and the presence of expired medicines and medicines not licensed for use in dairy cattle was also common. The medicine resources available to farmers are likely to influence their treatment decisions, therefore evidence of the PVM stored on farms can help inform understanding of medicine use.

## Introduction

Prescription veterinary medicine (PVM) use in the United Kingdom is an area of increasing focus for the veterinary profession, agricultural sector, government and food retailers (1). The agricultural sector is a significant user of antimicrobials (2), and reducing its overall use along with improving data collection were key recommendations of the O’Neill Report (3). According to the list published by the Veterinary Medicines Directorate (VMD) in the UK, as of 1^st^ January 2018 there are 1876 prescription-only veterinary medicines (POM-V) of which 142 are licensed to treat solely cattle and 456 are licensed to treat multiple species including cattle. These comprise 152 different listed active ingredients (4). There are 233 different antimicrobial preparations (containing 60 listed active ingredients) licensed for use in cattle with 75 licensed POM-V classed by the European Medicines Agency as highest-priority critically important antimicrobials (HP-CIA) to human health (5). Antimicrobial resistance is a recognised global threat and there have been many calls for improved understanding of antimicrobial prescription, use and recording (1, 3, 6-8). Currently in the UK, antimicrobial use (AMU) in the dairy industry is measured through a combination of pharmaceutical production/import data and more recently using a sample of individual veterinary practice sales data. This has improved data granularity but may not represent actual use on farms (2). Data for other PVM such as vaccines, non-steroidal anti-inflammatories (NSAIDs) or mechanical teat-sealants are not currently measured or published nationally.

Farmers in the UK can purchase and store PVM on farm for use at a later date (9). Many studies measure AMU on dairy farms (7, 10-15), however few/none report the quantity of antimicrobials stored on farms directly nor the way in which they are stored. Medicine storage on farms is an important part of compliance with Health and Safety Executive and farm assurance guidelines (16, 17), which require PVM to be placed in a secure, lockable location away from children, animals and thieves (18). In addition, medicines should be stored at the temperature requirements stated on the packaging. Despite this, a recent study found that vaccines were routinely being stored at inappropriate temperatures on UK farms (19). There is currently little evidence available to determine whether PVM are used in the way the prescribing veterinarian intended, or whether farmers are making decisions based on other factors while using stored PVM which may be expired or not licensed for use in dairy cattle.

There is evidence in human health research that prescription medicines are often stored in the home for use at a later date, despite the prescription having been intended for immediate use (20, 21). Studies have found that a proportion of patients deliberately planned to stop taking a course of prescribed antibiotics early in order to have a supply for self-use in the future (22, 23). Non-compliant use of medicines is commonly seen (24) and there is evidence that medicines are taken in ways other than that indicated by the prescriber (25, 26). It follows therefore that there is an urgent need for research into PVM storage and compliance in agriculture.

The aim of this study was to provide data on the storage practices of PVM on UK dairy farms and to investigate the quantity and composition of PVM being stored.

## Materials & Methods

### Study design & population

This research was approved by the University of Bristol Faculty of Health Sciences Research Ethics Committee under Approval No. 33021. This article was written according to the STROBE statement for scientific reporting of cross-sectional studies (27).

Data in this study formed part of a wider project investigating PVM use on UK dairy farms. Twenty-seven dairy farms in the South West of England and South Wales were enrolled. Veterinary practices within the study area were asked to recruit dairy farms. Mixed-species farms were only included where PVM purchase and storage for the dairy herd was kept separate from other PVM. Selection of farms was purposive based on varying herd size, production levels and management practices. Veterinary practices were asked to nominate farms across the spectrum of perceived medicine storage compliance to minimise selection bias. All farms were visited once by the lead author in a 6-week period in Autumn 2016.

On the day of the visit, a structured interview was conducted with the self-identified “main treatment decision-maker” (hereafter called the farmer) to gather information on farm demographics, management practices, animal health and productivity. Stock numbers, production, health and fertility data were ascertained to the best of the farmer’s knowledge aided by consultation with on-farm records. For the PVM inventory the farmer was asked to indicate areas on the farm where PVM might be found. The designated medicine cupboard was examined first, and certain high-probability storage areas were directly enquired about (e.g. household refrigerator, calf shed, milking parlour). Permission was also requested to search for PVM anywhere on the farm. A photograph of the medicine cupboard was taken and fieldnotes written as an *aide memoire* about the storage systems.

All PVM found were entered on-location into a pre-prepared spreadsheet. Location, drug name, pack size, number of packs, quantity remaining in each pack and expiry dates were noted. Where the product label was illegible it was disregarded. Where the expiry date was illegible it was assumed to be within date. Volume remaining was estimated by eye to the nearest 10% of pack size (i.e. for a 100 ml pack of liquid, volume was estimated to the nearest 10 ml and for a 50 g pack of powder, quantity was estimated to the nearest 5 g). All POM-V medicines were recorded along with any vaccines licensed for use in cattle and all pour-on, oral and injectable endectocides (anthelmintics). Vaccines were recorded in number of doses rather than volume. All intramammary and ocular medicines were recorded as single units per tube because one tube is equivalent to one dose.

### Data analysis

All data were entered into separate spreadsheets, one per study farm. The data from these spreadsheets were then collated and analysed by retrieving specific datasets through R software (28) and producing a database for all 27 farms to be compared and analysed together.

Medicine quantities were measured in total mg of active ingredient present, in mg per population corrected unit (PCU) in-line with nationally reported use data (29) and in total number of “medicine units” present. Medicine units were defined as: one bottle of liquid, one tube of intramammary or ocular suspension, one pack of boluses or tablets, one container of powder or one tube of ointment. Prescription veterinary medicines were grouped according to therapeutic group (e.g. antimicrobial, vaccine). Antimicrobials were grouped by antimicrobial class (e.g. penicillin, fluoroquinolone) and according to route of administration (e.g. injectable, intramammary). Expired medicines were defined as those medicines with an expiry date prior to the day of the inventory. Milligrams of active ingredient were presented to the nearest 100 mg for the total weight of active ingredient, to the nearest 10 mg for mg per cow in milk and mg per 1000 litres of milk produced annually and to the nearest 0.01 mg for mg/PCU.

Data was visually checked for normality. Normally distributed data were reported as a mean with standard deviation in brackets. Non-normally distributed data were reported as a median with range in brackets. Calculations were performed using a combination of Microsoft Excel (2016) and R (28). Given the cross-sectional, point-prevalence nature of the dataset and the fact that the study farms are not intended to be a representative sample of the population, presented calculations were descriptive and no inferences on causality can be made.

## Results

### Farm Demographics

Thirty-four dairy farms were identified as eligible for the study through self-nomination or nomination by veterinary practices. All were invited to enrol in the study, and 29 agreed to take part. Two farms dropped out of the study before the medicine inventory visit. Data for the remaining 27 farms was complete. These farms were located across seven counties and under the care of nine veterinary practices.

Farm demographic and management characteristics are described in Table 1. In summary, the median total herd size was 320 with 175 cows in milk. Most farms (59%) described the main cattle breed as Holstein and the majority calved year-round (81%). The median total annual milk volume per herd produced was 1.1 million L with annual milk sales per cow of 7500 L. Seventy-four percent of farmers had some formal specialised education and training in agriculture.

**Table 1:** Demographic and management characteristics of the 27 participating farms

| Characteristic | Response | Number of farms (n) |
| --- | --- | --- |
| Age of farmer (years) | 18-30 | 5 |
|  | 31-40 | 5 |
|  | 41-50 | 5 |
|  | >50 | 12 |
| Education level of farmer | Basic schooling | 3 |
|  | O Level/GCSE/A Level | 4 |
|  | HNC/HND/NVQ | 16 |
|  | University bachelor's degree | 4 |
| Total herd size | 100-199 | 6 |
|  | 200-299 | 12 |
|  | 300-699 | 10 |
|  | >700 | 5 |
| Total number of cows in milk | 1-99 | 4 |
|  | 100-199 | 10 |
|  | 200-299 | 7 |
|  | >300 | 4 |
| Calving pattern | Year-round | 22 |
|  | Seasonal – Spring | 4 |
|  | Seasonal – Autumn | 1 |
| Primary cow type | Holstein | 16 |
|  | British Friesian | 4 |
|  | Channel Island | 3 |
|  | Crossbreed | 2 |
|  | Other | 2 |
| Waste-milk feeding | Yes – all calves | 13 |
|  | Yes – beef calves only | 7 |
|  | No | 7 |
| Dry cow antimicrobial therapy | Blanket therapy | 18 |
|  | Selective therapy | 9 |

*Definitions: Farmer = self-described main treatment decision-maker; Basic schooling = no qualifications gained; GCSE = General Certificate of Secondary Education; HNC = Higher National Certificate; HND = Higher National Diploma; NVQ = National Vocational Certificate; Blanket therapy = all dry cows treated with intramammary antimicrobial; Selective therapy = certain dry cows not treated with intramammary antimicrobial based on somatic cell count and mastitis risk assessment*

Farm production and health characteristics are presented in Table 2. Two-thirds of farms used blanket dry cow therapy (where all cows were dried off with intramammary antimicrobial treatment). Twenty farms routinely fed waste milk containing antimicrobial residues to beef calves, 13 of which also fed waste milk to dairy replacement calves. The mean number of clinical cases of mastitis and lameness per 100 cows per year was 36.7 and 22.2 respectively. There were a median of 10 cases of respiratory disease and 10 cases of gastrointestinal disease per 100 calves per year.

**Table 2:** Production, health and medicine storage characteristics of the 27 participating farms

| **Property / characteristic** | **Median or Mean\*** | **Range (Median) or SD (Mean)\*** |
| --- | --- | --- |
| Total herd size | **320** | *119 – 1,271* |
| Number cows in milk | **175** | *65 – 600* |
| Total annual milk volume (million L) | **1.1** | *0.5 – 4* |
| Total annual milk sales per cow (L) | **7500** | *3,600 – 11,300* |
| Milk price (pence per L) | **22.5\*** | *15.3\** |
| Somatic cell count (cells per ml) | **176,407\*** | *50,466\** |
| Bactoscan (1,000 bacteria per ml) | **20.9\*** | *7.5\** |
| Mastitis (cases per 100 cows per year) | **36.7\*** | *15.9\** |
| Lameness (cases per 100 cows per year) | **22.2\*** | *15.3\** |
| Respiratory disease (cases per 100 calves per year) | **10** | *0 – 47* |
| GI disease (cases per 100 calves per year) | **10** | *3 – 59* |
| Number PVM present on farm (by active ingredient) | **19** | *9 – 35* |
| Number PVM present on farm (by medicine unit) | **101** | *28 – 339* |

*Median = non-normally distributed data Mean = normally distributed data*
*Definitions: PVM = Prescription veterinary medicines. L = Litre. SD = Standard deviation. GI = Gastrointestinal*

### Storage methods

Medicines were stored in six different location types across the study farms as seen in Figure 1. Most were stored in a lockable medicine cupboard or refrigerator although 29% were stored in a non-compliant area such as the milking parlour, the calf shed or the office. Ten farms stored 100% of their medicines in lockable medicine cupboards or lockable refrigerators, and two farms did not store any medicines in a lockable medicine cupboard or lockable refrigerator. No participating farm monitored the temperature of their refrigerator or medicine storage area.

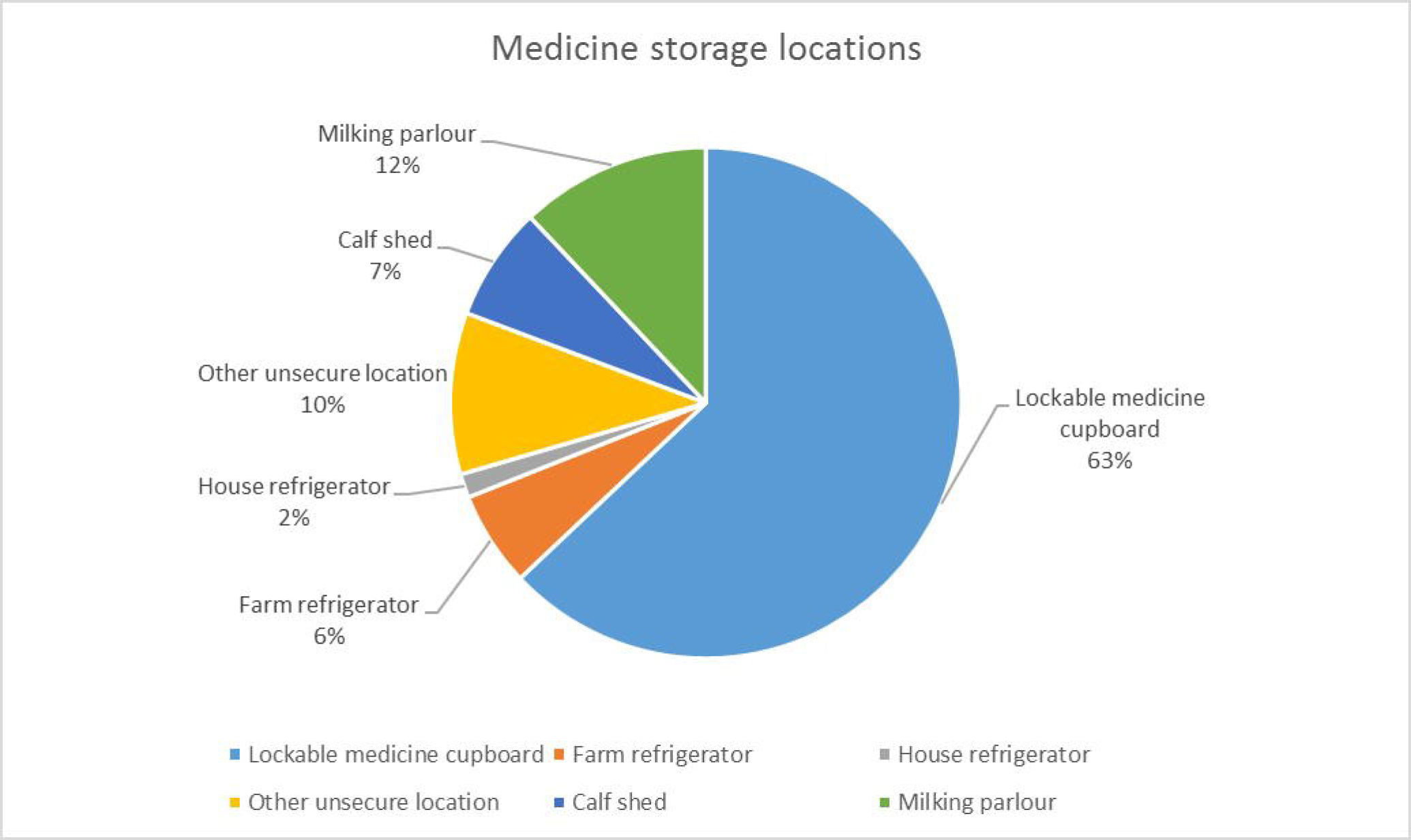

### Prescription Veterinary Medicines

There were a median of 19 (9-35) different types (by active ingredient) of PVM and 101 (28-339) individual medicine units of PVM present on participating farms. Antimicrobials were the therapeutic group most commonly stored both by frequency of occurrence (median 69 (22-296) medicine units) and by total weight (median 182,300 (45,500 – 442,500) mg) equivalent to 1.54 mg/PCU.

#### Antimicrobials

The route of administration for antimicrobials stored are presented in Table 3. Of the total antimicrobials stored across all farms 76.4% were injectable and 14.7% were intramammary. When units were measured, 10% were injectable (bottles) and 84.8% were intramammary tubes.

**Table 3:** Total quantities of antimicrobial stored across all 27 participating farms, by route of administration

| Route of administration | Total (mg) | Total (medicine units) |
| --- | --- | --- |
| All | 5,127,176 (100%) | 2,801 (100%) |
| Injectable | 3,917,116 (76.4%) | 280 (10%) |
| Intramammary | 755,772 (14.7%) | 2,374 (84.8%) |
| Topical | 284,348 (5.5%) | 51 (1.8%) |
| Oral | 54,440 (1.1%) | 31 (1.1%) |
| Intrauterine | 115,500 (2.3%) | 65 (2.3%) |

Data on antimicrobial storage is presented in Table 4. Total HP-CIA storage per farm was a median of 10,000mg, or 0.12 mg/PCU with the majority (5000mg) being 3^rd^ generation cephalosporin. Injectable antimicrobials comprised the greatest total weight with a median of 143,600mg or 1.19 mg/PCU, compared with intramammary antimicrobials which were a median of 21,200mg or 0.21 mg/PCU. Conversely, intramammary antimicrobials had the greatest total number of medicine units present, with a median of 66 units compared with 9 units of injectable antimicrobials.

**Table 4:** Quantity of antimicrobial stored on the 27 participating farms

| Antimicrobial type | Total (mg) |  | Total (mg/PCU) |  | Total (Medicine units) |  | Number of farms where present |
| --- | --- | --- | --- | --- | --- | --- | --- |
|  | Median | Range | Median | Range | Median | Range |  |
| Total antimicrobial | 182,200 | 45,500 – 442,400 | 1.54 | 0.51 – 5.08 | 69 | 22 – 296 | 27 |
| Total HP-CIA | 10,000 | 0 – 68,000 | 0.12 | 0.00 – 0.34 | 5 | 0 – 95 | 24 |
| Fluoroquinolone | 0 | 0 – 32,000 | 0.00 | 0.00 – 0.19 | 0 | 0 – 4 | 11 |
| 3<sup>rd</sup> gen. cephalosporin | 5,000 | 0 – 36,000 | 0.06 | 0.00 – 0.30 | 1 | 0 – 5 | 19 |
| 4<sup>th</sup> gen. cephalosporin | 1,000 | 0 – 9,000 | 0.01 | 0.00 – 0.12 | 0 | 0 – 59 | 13 |
| Injectable (all) | 143,600 | 16,500 – 393,000 | 1.19 | 0.19 – 4.51 | 9 | 1 – 35 | 27 |
| Intramammary (all) | 21,200 | 1,000 – 111,400 | 0.21 | 0.01 – 0.49 | 66 | 5 – 248 | 27 |
| Other (all) | 13,200 | 0 – 61,800 | 0.11 | 0.00 – 0.63 | 5 | 0 – 24 | 21 |
| Injectable (HP-CIA) | 5,000 | 0 – 36,000 | 0.09 | 0.00 – 0.34 | 2 | 0 – 6 | 24 |
| Intramammary (HP-CIA) | 1,000 | 0 – 9,000 | 0.01 | 0.00 – 0.12 | 0 | 0 – 93 | 12 |
| Other (HP-CIA) | 0 | 0 | 0.00 | 0.00 | 0 | 0 | 0 |

*Definitions: HP-CIA = Highest Priority Critically Important Antimicrobial (Ref EMA); 3<sup>rd</sup> gen. cephalosporin = 3<sup>rd</sup> generation cephalosporin; 4<sup>th</sup> gen. cephalosporin = 4<sup>th</sup> generation cephalosporin*

Eighty-nine percent of farms stored at least one HP-CIA. The most frequently occurring injectable antimicrobials were ceftiofur (a HP-CIA; n=24) and penicillin/streptomycin combination (n=24). Also commonly found were oxytetracycline (n=22), tylosin (n=19) and trimethoprim/sulfadiazine (n=16). The three lactating-cow intramammary antimicrobials most commonly identified were potentiated amoxycillin (n=11), the combination streptomycin/neomycin/novobiocin/penicillin (n=11) and cefalexin/kanamycin (n=10). The most frequently occurring dry cow intramammary antimicrobials were cephalonium (n=12), cefquinome (a HP-CIA; n=10) and cloxacillin (n=8).

The total mg/PCU on each farm ranged from 0.51 - 5.08 mg/PCU (Figure 2). Total mg per cow in milk ranged from 430 – 3,430 mg (Figure 3). Total mg per 1000 litres of milk produced annually ranged from 40 – 740 mg (Figure 4).

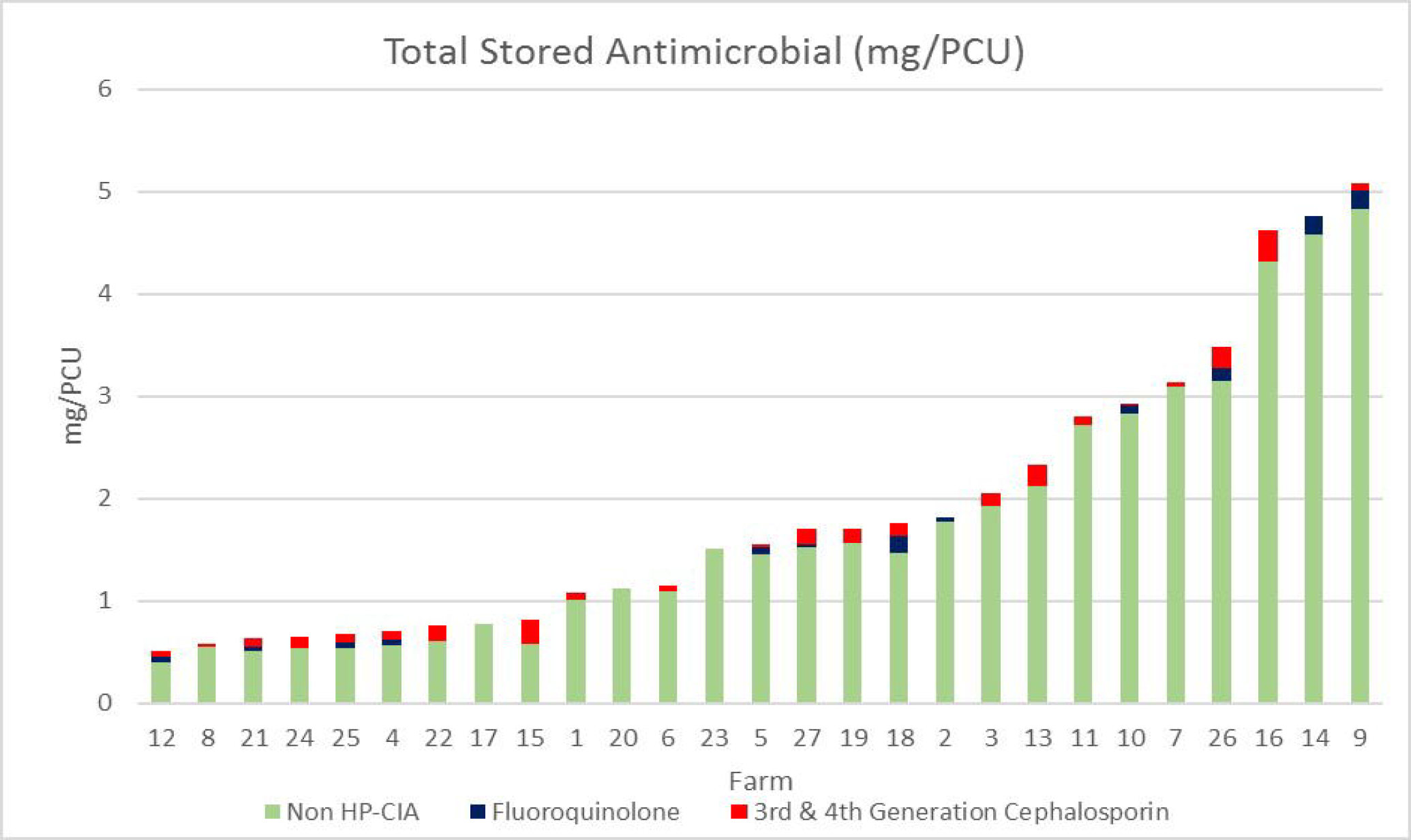

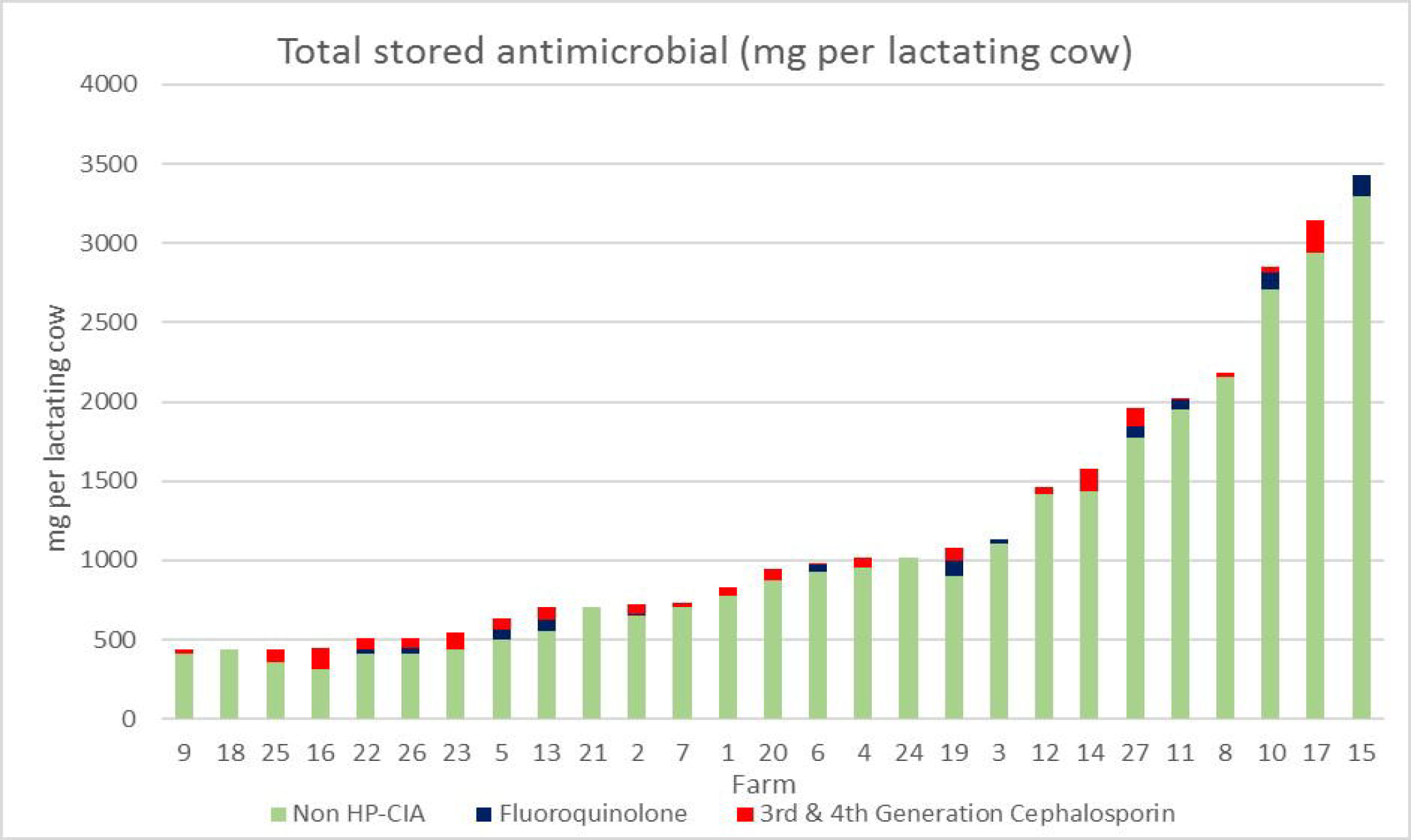

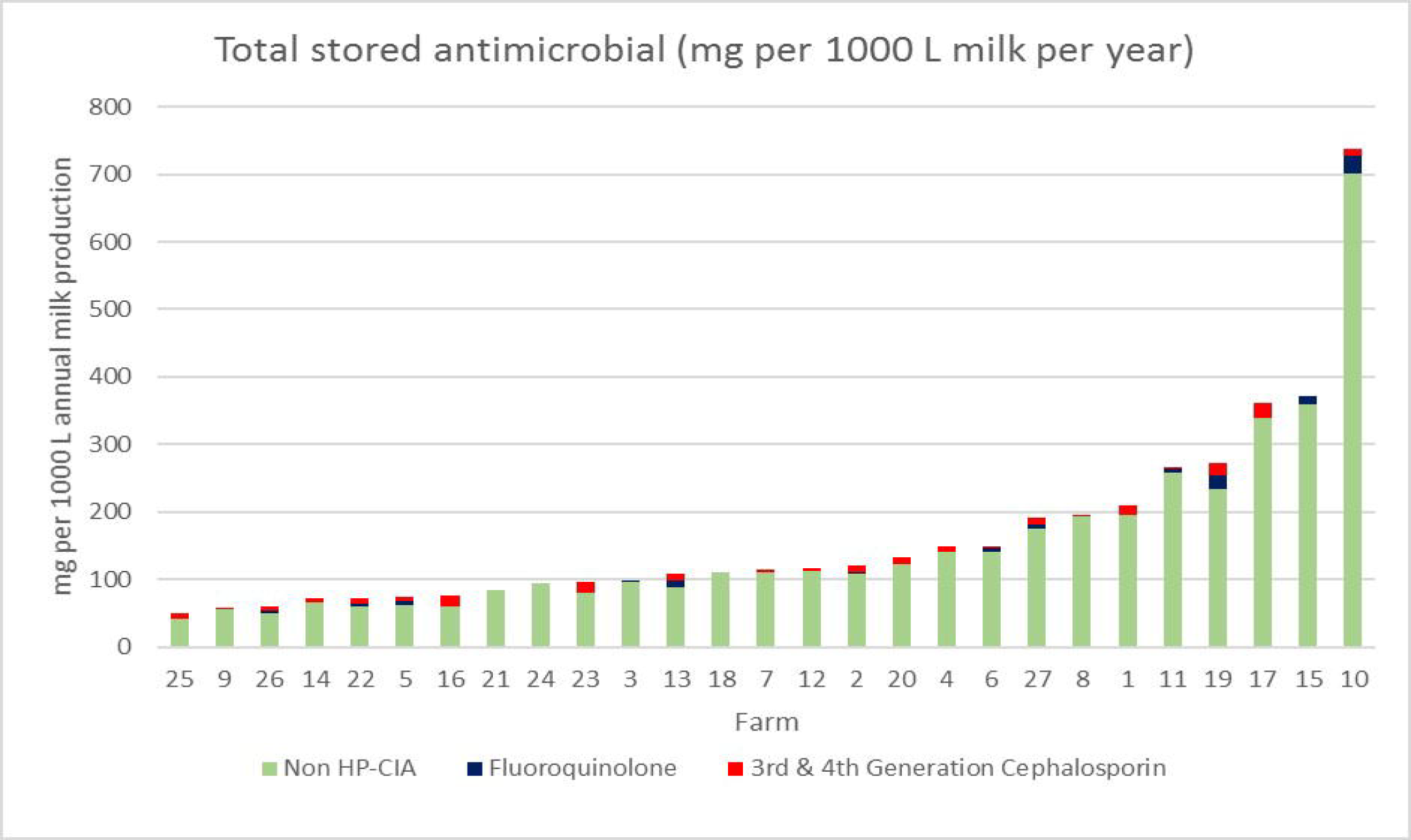

#### Vaccines & Other PVM

The total number of vaccine doses stored across all farms was 3,541 with a median of 0 (0-1893) doses per farm. Eighteen farms (66.7%) stored no vaccines at the time of the study. The most common diseases for which vaccines were stored were Bovine Herpes Virus-1 (866 doses; 5 farms), leptospirosis (835 doses; 5 farms) and bovine viral diarrhoea (BVD) (683 doses; 5 farms). Other diseases for which vaccines were stored were calf diarrhoea, bluetongue, clostridial diseases, lungworm, mastitis and calf pneumonia.

The total number of units of NSAIDs across all farms was 75, with a median of two per farm (0-8). The total number of anthelmintics or anti-protozoals present across all farms was 87, also with a median of 2 per farm (0-14). The total number of units of hormone across all farms was 53, with a median of 1 (0-10) unit per farm. Five farms (19%) stored prostaglandins.

### Expired medicines

At least one unit of expired PVM was stored on 25 farms (93%). Eighteen farms (67%) stored at least one expired antimicrobial and six (22%) stored at least one expired vaccine. The total number of expired antimicrobial units across all farms was 201, with each farm storing a median of 2 (0-58) expired antimicrobials. The total number of doses of expired vaccines stored across all farms was 827, with each farm storing a median of 0 (0-725) expired doses.

The median length of time since expiration was 12 (1-200) months. For antimicrobials, median length of time since expiration was 10 (1-96) months, for anti-inflammatories 6 (2-46) months, for vaccines 20 (2-200) and for other medicines 12 (2-120) months, with the majority of the “other” medicines endectocides.

### Cascade medicines

Medicines not licensed for use in dairy cattle were found on 16 farms (59%). A total of 30 unlicensed units were stored comprising 14 different medicines. Seven different unlicensed antimicrobials were identified totalling 709,100 mg of active ingredient. Macrolides were most common with 138,600 mg of erythromycin (licensed for use in poultry and pigs) found across seven farms served by two different veterinary practices - 2,000 mg lincomycin on one farm (licensed for use in pigs) and 200 mg gentamycin (licensed for use in horses not producing milk or meat for human consumption) on one farm. Also found were 500,000 mg of tetracyclines (licensed for use in poultry and pigs) across three farms served by one veterinary practice, 5,000 mg metronidazole veterinary tablets (licensed for use in dogs and cats but banned from use in cattle) on one farm, 18,000 mg of 1st generation cephalosporin of Hungarian origin and not licensed for sale in the UK on one farm and 30,000 mg of florfenicol of Dutch origin not licensed for sale in the UK on a different farm. There was no clustering of unlicensed products by farm. Other unlicensed medicines identified were: 50 ml of mepivacaine (a local anaesthetic agent licensed for use in horses), 1 tube of acepromazine gel and 30 ml romifidine (sedatives licensed for use in horses), 2 capsules of 2 mg loperamide (an antispasmodic licensed for human use), 200 ml sucralfate, 2 tubes of fusidic acid gel (an antibiotic topical gel licensed for use in cats and dogs) and one bottle of topical miconazole licensed for use in dogs. Where unlicensed medicines were found, farmers confirmed they were present for use in the dairy calves or adult cattle.

## Discussion

Most farms in this study stored PVM in the recommended way. However, some PVM were being stored in inappropriate conditions e.g. in areas at risk from heat or cold damage and exposure to sunlight or gross contamination. Twenty-nine percent of PVM found were not stored in a lockable cupboard or room. This has direct health and safety implications due to access to potentially harmful medicines by animals or children in addition to risks of theft. Certain medicines should be stored with particular care due to their potential for harm from accidental exposure (18). Prostaglandins for example, which made up a proportion of the hormone POM-V reported in the results section, can be absorbed transcutaneously and lead to miscarriage or serious and even fatal respiratory compromise in susceptible people (30).

While some farms stored a wide range of different types and quantities of PVM, others stored a limited number. The fact that the quantity of antimicrobials stored on farm does not appear to be linked with the number of animals at risk of treatment or the overall production values of the cows on the farm (Figures 2-4) suggests that there are other reasons for the range of storage practices seen. Ongoing work by the lead author, as part of a wider study, is exploring these reasons (31).

As previously noted, a farmer’s treatment decisions are to some extent constrained by the PVM resources available to them. It follows therefore that when designing policy interventions aimed at reducing AMU, data on storage practices and farmers’ use of stored medicines is extremely important.

### Antimicrobial storage

Antimicrobials were the PVM stored in the greatest quantity when measured by total mg of active ingredient as well as by individual medicine units. Twenty-four farms stored HP-CIAs indicating that their use is still common in UK dairy farming, however recent increased efforts to reduce their use mean this is likely to be a rapidly evolving picture. For example, as of June 2018 Red Tractor Farm Assurance will require HP-CIAs only be used as a last resort, with a veterinary report outlining diagnostic or sensitivity testing (32). While storage does not equate to actual use, it is likely that these antimicrobials are stored with the intention of use and therefore it may be that the use of HP-CIAs was still common practice during the period of the study. Data from the wider longitudinal research study is in preparation for publication and will report on actual PVM use on these farms including HP-CIA use over a 12-month period.

The number of bottles of injectable antimicrobial present on farms was as high as 35 on one farm, with a median of 9. Keeping a store of antimicrobials like this provides a large resource for the farmer to use without a need to consult her/his veterinarian. One of the most frequently kept injectable antimicrobials was ceftiofur, a HP-CIA. Given the focus on reducing the use of HP-CIAs in the years preceding the study, this may be indicative of a reluctance to move away from their use in the dairy sector. The 0-hour milk withhold carried by ceftiofur and it’s broad licensing for use in respiratory disease, metritis and interdigital necrobacillosis made it an attractive and cost-effective option for treating disease on farm, and it appears to have remained popular at least until the latter part of 2016. The antimicrobials most commonly used for treating mastitis in the lactation period were not on the current HP-CIA list although potentiated amoxycillin is not considered to be a first-line treatment (33). The most commonly stored antimicrobials for treating dry cows included cefquinome, a 4^th^ generation cephalosporin and HP-CIA.

Many of the first-line, “responsible” antimicrobials have a relatively high total weight of active ingredient when compared with HP-CIAs, leading to calls for HP-CIAs to be measured and benchmarked separately from other antimicrobials (6, 7). This study provides evidence that there in an ongoing need to change behaviour and reduce the use of HP-CIAs on dairy farms.

Interestingly, when measured in mg/PCU or mg/1000L milk produced annually, the data show that while most farms stored similar quantities of antimicrobial, a handful of farms stored up to ten times as much as those farms which stored the smallest amounts. This suggests other factors affect the storage practices of dairy farmers, something being investigated in ongoing work by the authors.

Participating farms stored a broad range of different antimicrobials, thus increasing their options when making treatment decisions. This could lead to a dissonance between the intention of the prescribing veterinarian and the actions of the farmer. Having such a large resource to draw upon could be seen to improve the agency and ownership of the farmer on those decisions, but conversely to decrease the agency and ownership of the veterinarian legally responsible for their use. This serves to emphasise the importance of understanding the treatment decisions, given the relatively few resource constraints.

### Expired medicines

While the presence of expired medicines does not equate to their use, the fact that expired PVM were identified on most participating farms indicates that their use is likely to be common. Expiry dates for drug products are set based on real-time stability testing at appropriate storage conditions to determine whether the drug substance meets its individually set specification (34). A specification “establishes the set of criteria to which a drug product should conform to be acceptable for its intended use” (35). All but two farms stored at least one expired PVM with two-thirds storing at least one expired antimicrobial. This is particularly striking when compared with studies of household medicine storage among human health, which have shown a range of 3-22% of stored medicines were expired (36, 37). Given the average length of time passed since expiry was 12 months, with one farm storing medicine that was over 16 years out-of-date, their presence appears to be accepted by farmers on dairy farms.

The impact of using an expired antimicrobial is ill-defined. It is assumed that the efficacy of an antimicrobial, or indeed any medicine, reduces with time after expiration. However, the evidence base for this is small and contradictory. In one study from human health, it was shown that there was a decreased rate of pathogen susceptibility to expired antimicrobials (38). Other studies have shown that most medicines retain their efficacy for many years beyond their expiration date (39, 40). To the authors’ knowledge there are no studies on the efficacy of expired antimicrobials in veterinary medicine.

Perhaps more important than the expiry date stated on PVM is the shelf life of the medicine once broached. In-use shelf life is determined for multi-use veterinary products by in-use stability testing of physical, chemical and microbial properties. Products approaching the end of their shelf life are tested, with testing designed to simulate as closely as possible real-life conditions based on likely usage patterns of the product under “normal environmental conditions” and stored according to the product literature. These drugs are measured against either their original specification or an “in-use shelf life” specification, as appropriate (41). While this is often 28 days for injectable products and 24 hours for vaccines, in reality these shelf lives are rarely observed due to most injectable medicines being sold in 100 or 250 ml multi-dose bottles and individual animals’ treatment courses require varying volumes of medicine. Measuring the presence and use of PVM that had passed its broached shelf-life was beyond the scope of this study, however future research in this area would be valuable.

Expired or waste PVM should not be disposed of with normal household waste and most veterinary practices offer a disposal service to clients. Given the prevalence of expired PVM on the study farms, veterinarians should determine whether the farms under their care are disposing of these medicines appropriately or whether they remain on farm with potential for use. Discussion of the use and disposal of expired PVM would make a valuable addition to herd health review meetings, particularly given the veterinarian’s ultimate responsibility for the safety of medicines being used in these food-producing animals.

### The Cascade

Using PVM which are not licensed for use in dairy cattle is not illegal if they are prescribed and used according to the Cascade and where there are established Maximum Residue Limits (42). However, the use of unlicensed PVM is not currently monitored in the UK. Given the presence of medicines which have been prescribed via the Cascade further research is urgently needed in this area. In one instance, PVM were present which are explicitly banned from use in dairy cattle (metronidazole): administration would constitute a transgression of the law (43).

### Study Limitations

The use of purposive sampling through veterinary practice nomination inevitably leads to the possibility of selection bias. The study farms were demographically reflective of the wider UK dairy farm population. According to the AHDB the “average number of adult dairy cows” on UK dairy farms in 2016 was 143 (44), compared to the study farm median of 175. The larger herd size of the study farms may influence the way in which PVM are stored. Larger herds are more likely to have an increased frequency of veterinary visits which may mean they store fewer PVM on the farm as they have additional resources available to them through the veterinarian on a regular basis.

Farmers were asked not to alter the medicines stored on their farms for the visit day. Given the prevalence of expired medicines and storage of medicines outside of designated cupboards it appears that farmers did not significantly improve their storage practices prior to the visit, and the authors believe the data described are representative of normal medicine storage on the study farms. The volume of medicines in opened bottles was estimated by eye to the nearest 10 ml, which may have led to some over- or under-estimation of the total volume present on farm. Using reference weights for different medicines and a portable weighing scale to measure the weight of bottles may have improved the accuracy of these measurements.

This study was cross-sectional, and the seasonal nature of dairy farming and disease prevalence should be noted. This study took place in the Autumn, around the time of housing for many farms, and the data may be different if it was to be repeated in different seasons. Where mixed-enterprise farms were included, these stored medicines intended for use in dairy cattle and calves separately from medicines intended for beef or sheep. This did not allow for any measurement of the possibility of medicines stored for use in beef and sheep being used in the dairy cattle. It is also important to note that this study reports storage practices on a small number of dairy farms in South West England and South Wales and as such may not reflect practices found on other farms or in other regions of the UK. Further research in this area is needed to provide a robust evidence base for future policy decisions aimed at improving responsible medicine use in dairy farming.

### Conclusions

These are the first data of their kind published on the UK situation and are useful to help veterinarians understand the ways in which medicines are being used post-prescription and to inform future herd health planning. Current UK estimates of PVM use are crude and only through detailed on-site research can real medicine use practices be discerned. The results are also helpful for policy makers and researchers to broaden the evidence base surrounding PVM use.

## Acknowledgements

The authors would like to thank the participating farmers and veterinary practices for their help with this study.

